# Circulating miR-10b-5p drives Kawasaki vasculitis through endothelial reprogramming and nominates CXCL8 as an early diagnostic biomarker

**DOI:** 10.64898/2026.03.12.711476

**Authors:** Sunyoung Park, Gaeun Kang, Mira Kim, Min Hee Kim, HyeonJin Yang, Bitna Lim, Seunghoon Jung, Young Kuk Cho, Young-Kook Kim, Woo-Jae Park, Somy Yoon, Gwang Hyeon Eom

## Abstract

**Background:** Kawasaki disease (KD) is an acute systemic vasculitis of unknown etiology and the leading cause of acquired heart disease in children. Coronary artery aneurysm formation represents its most serious complication, yet no specific biomarker exists for early diagnosis at emergency department (ED) presentation. We sought to define the molecular mechanisms driving KD vasculitis and to identify a clinically actionable early diagnostic marker.

**Methods:** Whole microRNA sequencing was performed on buffy coat specimens from KD patients and febrile controls presenting to the ED. Disease-associated miRNA candidates were validated in two independent murine KD-like vasculitis models: *Lactobacillus casei* cell wall extract (LCWE; n=12–20) and *Candida albicans* water-soluble fraction (CAWS; n=12). The functional role of miR-10b-5p was assessed by antagomir-mediated inhibition in the LCWE model. Transcriptomic and chromatin-level reprogramming were characterized by single-cell RNA sequencing (scRNA-seq), bulk RNA-seq, and assay for transposase-accessible chromatin with sequencing (ATAC-seq) in human coronary artery endothelial cells (HCAECs). Transcription factor binding was validated by *in vitro* binding assays and chromatin immunoprecipitation. Serum chemokine (C-X-C motif) ligand 8 (CXCL8) was quantified by enzyme-linked immunosorbent assay.

**Results:** Forty miRNAs were upregulated in KD patients and fourteen miRNAs were prioritized for further analysis. miR-10b-5p was consistently elevated across both vasculitis models, and its antagomir-mediated inhibition attenuated aortic root inflammation *in vivo*. In HCAECs, miR-10b-5p suppressed its direct targets *marker of proliferation Ki-67 (MKI67)*/*Chromobox5 (CBX5),* driving a cell-state transition from proliferative/metabolic programs toward a pro-inflammatory phenotype. Consistently, analyses of bulk RNA-seq supported cell-cycle arrest and altered chromatin remodeling. ATAC-seq revealed increased chromatin accessibility at the *CXCL8* and *Matrix metalloproteinases 10* (*MMP10)* promoters, and motif analysis identified CCAAT/enhancer-binding protein alpha (CEBPA) as the key transcriptional activator; *CEBPA* knockdown abrogated miR-10b-5p–induced *CXCL8* and *MMP10* upregulation. Serum CXCL8 was markedly elevated in KD patients relative to febrile controls at first ED presentation, prior to definitive diagnosis, and declined significantly at 8-week follow-up.

**Conclusions:** We define a miR-10b-5p–MKI67*/*CBX5–CEBPA–CXCL8 axis as a mechanistic driver of endothelial reprogramming and neutrophil-recruiting inflammation in KD. Serum CXCL8 emerges as a candidate early diagnostic biomarker that may facilitate timely KD recognition at ED presentation.

## Introduction

Kawasaki disease (KD) is an acute systemic vasculitis that primarily affects in young children in northeast Asia^1–3^, with 85% of patients under 5 years of age^4–6^. It is characterized by persistent fever with mucocutaneous inflammation and lymphadenopathy, and its most serious complication is coronary artery involvement, including coronary artery aneurysms. Despite extensive research aimed at elucidating the pathophysiology of KD, its etiology and underlying mechanisms remain unclear^1,2,7^. However, cumulative evidence has shown that KD is related to synergistic outcome of genetic susceptibility and excessive immune response^1,2,5^. Geographically, KD is prone to occur in northeast Asian populations^1–3^, which suggests a genetic predisposition to KD. The most common and distinctive clinical feature of KD is a persistent high fever that does not resolve with anti-inflammatory medications. Thus, infection is considered another contributing factor to cause KD^1,2^. It is widely accepted that KD may occur when genetically susceptible people are exposed to infectious agents, such as bacteria and viruses^5^. The most serious and potentially fatal complication of KD is coronary artery aneurysm formation^4,6^. In fact, coronary artery aneurysms caused by KD can be progressive and are considered a leading cause of mortality from acquired heart disease in childhood and young adulthood^8^. Hence, early diagnosis and treatment are crucial to minimize the risk of coronary aneurysm in KD^4,6^. Still, intravenous immunoglobulin (IVIG) is the treatment of choice^9^, but it is known that the timing of first intervention for treatment can determine the disease prognosis^4,6^. In other words, it is highly important to evaluate the possibility of KD when a patient first visits the clinic. IVIG treatment alleviates acute symptoms of KD in many cases; however, the proportion of IVIG-refractory KD—defined as persistent or recurrent fever after the initial IVIG infusion—appears to be increasing^10,11^. Accordingly, there remains an unmet medical need to elucidate the pathogenesis of KD and to develop complementary strategies to improve its diagnosis and treatment.

Currently, there are no specific diagnostic laboratory markers for KD^4,6^. For diagnosing KD, diagnostic criteria for complete KD and suspected incomplete KD are used^4,6^. The signs and symptoms include fever, rash, conjunctivitis, oral changes, erythema, and cervical adenopathy^4,6^. They start with fever or rash^4,6^, which are nonspecific from other infectious diseases. Fulfilling these diagnostic criteria generally takes time, which can delay the diagnosis and intervention of KD^12^. This issue is particularly challenging in incomplete KD, which presents with prolonged fever but fewer of the classic clinical features^12^.

Regarding KD vasculitis, smooth muscle cells and endothelial cells are both considered the key contributors of vascular inflammation, and overall, the endothelium-centered theory is broadly supported these days^13^. Vascular endothelial cells are directly exposed to circulating cells and pathogens within the vessel lumen and are well recognized as key regulators of immune responses. By secreting chemotactic factors and engaging in early interactions with immune cells (i.e., neutrophils), endothelial cells can initiate, amplify, or terminate inflammatory responses^14^. During the development of coronary artery lesions, extracellular matrix (ECM) breakdown and consequent weakening of the vascular wall occur^15^. Therefore, ECM-degrading proteases, such as matrix metalloproteinases (MMPs), have emerged as key mediators of this process^16^. Although the pathophysiology of KD has not yet been clearly elucidated, it is generally accepted that neutrophils play a crucial role in the hyperacute phase of KD^7,17,18^. As initial neutrophil activation leads to the following necrotizing and chronic vasculitis^7^, thoroughly investigating how neutrophils are activated is necessary.

In KD, vasculitis often persists even after the acute clinical symptoms have resolved^4,6^. Coronary artery lesions can continue to progress despite the absence of ongoing KD or overt infectious diseases^19^. This pattern suggests that one or more factors capable of sustaining vascular inflammation remain present within the vasculature after the initial disease course. Given their relative stability and potential for long-term persistence in circulation^20^, microRNAs (miRNAs) are plausible candidates that may contribute to prolonged stimulation of KD vasculitis. miRNAs were first discovered in KD patients’ whole blood in 2013^21^, although their functions were not investigated enough. It was also reported that miRNAs affect vascular endothelial cells in KD^22,23^.

Here we hypothesized that dysregulated miRNAs contribute to KD vasculitis. To test this, we first profiled circulating miRNAs in patient blood by miRNA sequencing and then evaluated the functional role of miR-10b-5p in an *in vivo* KD model and in a human coronary artery endothelial cell (HCAEC)–based model using transcriptomic and chromatin-accessibility analyses. We found that miR-10b-5p induces cell-cycle arrest and chromatin remodeling, leading to increased CXCL8 secretion. Importantly, CXCL8 was elevated at high levels from a very early stage—before KD could be definitively diagnosed—highlighting its potential diagnostic value.

## Methods

### Human samples

Febrile control and KD patient blood samples were collected in Chonnam National University Hospital from 2015 to 2017 under the Chonnam National University Institutional Review Board (CNU IRB, No. I-1009-09-103). Patients presenting to the ED with acute febrile illness were identified as the source population. All eligible ED visits underwent routine clinical evaluation and diagnostic work-up according to standard care. Based on the final diagnosis established after completion of the evaluation, patients were retrospectively classified into two groups: (1) a febrile control group, comprising patients in whom a specific non–KD etiology for fever was identified; and (2) a KD group, comprising patients with a final confirmed diagnosis of KD. KD was defined according to established diagnostic criteria.

Peripheral blood samples were collected at the initial ED visit and again at the 8-week follow-up outpatient visit after hospital discharge. Blood samples were incubated at room temperature (RT) for 30 minutes and centrifuged at 4 ℃ at 3,000 revolutions per minute (RPM) for 20 minutes. Patient sera were collected and stored at -80 ℃ before analysis. The use of archived human samples was reviewed and approved separately (CNU IRB, No. CNUH-2023-177).

### Cell culture

Primarily cultured human coronary artery endothelial cells (HCAECs) were purchased from Lonza (Basel, Switzerland) and cultured with EGM-2MV Microvascular Endothelial Cell Growth Medium-2 BulletKit (Lonza, Basel, Switzerland) on a 60 mm dish with 5% CO2 at 37 ℃. The complete growth medium was stored at 4 ℃ and used within 1 month. Cells were subcultured when they were 70-85% confluent and plated with 30-50% confluency. The passage number was kept under 10. We used five different batches of HCAECs in this study: 22TL108498, manufactured on July 8, 2022, from a 36-year-old female donor; 23TL363491, manufactured on December 31, 2023, from a 37-year-old female donor; 24TL031385, manufactured on March 3, 2024, from a 28-year-old male donor; 24TL065710, manufactured on April 8, 2024, from a 64-year-old male donor; and 24TL151939, manufactured on October 4, 2024, from a 48-year-old female donor. Biochemical and phenotypic responses were consistent across batches.

Primarily cultured human coronary artery smooth muscle cells (CASMCs) were purchased from Lonza (Basel, Switzerland) and cultured with Clonetics SmGM-2 BulletKit (Lonza, Basel, Switzerland). We used 0000641247, manufactured on July 26, 2017, from a 56-year-old male donor.

HEK 293T cells were cultured with Dulbecco’s Modified Eagle Medium (DMEM High; HyClone, Logan, UT, USA) with 10% HyClone Characterized Fetal Bovine Serum (FBS; HyClone, Logan, UT, USA) and 1% HyClone Penicillin-Streptomycin Solution (HyClone, Logan, UT, USA) with 5% CO_2_ at 37 ℃.

### Lactobacillus casei cell wall extract

*Lactobacillus casei* cell wall extract (LCWE) was prepared from *Lactobacillus casei* (11578, ATCC, Manassas, VA, USA) as previously described, with minor modifications. Briefly, bacteria were cultured in MRS broth at 37 °C and harvested during the exponential growth phase by centrifugation. The pellet was treated overnight with 4% sodium dodecyl sulfate (SDS), followed by sequential incubations with RNase, DNase I, and trypsin (each at 250 µg/mL for two hours) to remove contaminating nucleic acids and proteins. SDS-treated cell-wall fragments were resuspended in PBS and sonicated for 2 h while maintaining the preparation cold using a dry ice/ethanol bath then clarified by centrifugation of 20,000 × relative centrifugal force (RCF) for 1 hour at 4 °C. The supernatant containing LCWE was collected, and LCWE concentration was determined based on rhamnose content using a colorimetric phenol–sulfuric acid assay and adjusted to the desired working concentration. Aliquots were stored until use. One milligram of LCWE or the same volume of PBS were i.p. injected in male mice aged 5 weeks old (BALB/c). Mice were purchased from Orient Bio (Seongnam, South Korea), and vehicle or LCWE was injected in a single-blinded manner. No mice were excluded in this study. All animal studies were reviewed and approved by the Institutional Animal Care and Use Committee (CNU IACUC-H-2017-31).

### Preparation of CAWS (*Candida albicans* water-soluble fraction)

CAWS was prepared from *Candida albicans* (NBRC 1385) following a published protocol. Briefly, C. albicans was cultured in a complete synthetic, carbon-limiting medium (pH 5.2) at 25 °C with aeration (room air) and continuous agitation of 400 rpm for 2 days. The culture supernatant was then mixed with an equal volume of ethanol and incubated overnight to precipitate ethanol-insoluble material. The precipitate was collected, extracted with distilled water to recover the water-soluble fraction, and re-precipitated by adding ethanol followed by overnight incubation. The final precipitate was collected and dried with acetone to obtain CAWS, which was subsequently reconstituted in PBS, aliquoted, and stored at −80 °C until use. KD–like vasculitis was induced by intraperitoneal injection of CAWS (4 mg per mouse, BALB/c) for 4 weeks. CAWS models were generated in a single-blinded manner.

### Quantification of circulating miRNAs

Whole blood was collected from LCWE or CAWS-treated mice at 4 weeks after administration. Total RNA, including small RNAs, was extracted directly from whole blood using TRIzol (Invitrogen, Waltham, MA, USA). miRNA quantification was performed using the miRCURY miRNA PCR system (QIAGEN, Venlo, the Netherlands) following the manufacturer’s protocol, including reverse transcription and quantitative polymerase chain reaction (RT-qPCR) with miRNA-specific assays. Relative expression of miRNA was determined using the ΔΔCt method normalized to U6 as the internal reference control.

### miRNA mimic treatment

*mir*Vana miRNA Mimic, Negative Control #1 (4464061, Invitrogen, Waltham, MA, USA) or *mir*Vana miRNA Mimic, hsa miR-10b-5p (UACCCUGUAGAACCGAAUUUGUG, 4464066, Invitrogen, Waltham, MA, USA) was reconstructed with Nuclease-Free Water (Invitrogen, Waltham, MA, USA) as 100 μM stock and transfected to HCAECs as final concentration of 30 nM using Lipofectamine RNAiMAX Transfection Reagent (Invitrogen, Waltham, MA, USA).

### Locked nucleic acid (LNA) miRNA antagomir

An *in vivo* rescue experiment was performed using an mmu-miR-10b-5p antagomir. A miRCURY LNA miRNA inhibitor targeting mouse miR-10b-5p (5′-TCGGTTCTACAGGGT-3′; 339203 YCI0203119-FZA), miR-125a-3p (5’-CAAGAACCTCACCTG-3’; 339203 yci0203118-FZA) and a matched negative control inhibitor (5′-ACGTCTATACGCCCA-3′; 339203 YCI0202524-FZA) were custom-designed and supplied by QIAGEN (Venlo, Netherlands). Both oligonucleotides contained LNA modifications to enhance *in vivo* stability; the exact positions of the LNA nucleotides were not disclosed by the manufacturer. One milligram of the antagomir was administered via tail-vein injection in a single-blinded manner, concurrently with LCWE administration. To account for its circulating half-life, a second dose was administered two weeks later. Inhibition efficacy was assessed using the miRCURY miRNA PCR system (Y00205689, QIAGEN, Venlo, Netherlands).

### Masson’s trichrome staining for aortic valve inflammation

Aortic roots including the aortic valve were harvested at 4 weeks after LCWE or LNA miRNA antagomir. Heart was rinsed in PBS, and fixed in 4% paraformaldehyde for overnight. Tissues were processed, paraffin-embedded, and sectioned at 5 µm. Masson’s trichrome staining was performed using a commercial kit (ab150686, Abcam, Cambridge, United Kingdom) according to the manufacturer’s instructions. Briefly, sections were deparaffinized, rehydrated, and subjected to sequential staining to visualize collagen deposition (blue) and muscle/cytoplasm (red). Stained sections were imaged using bright field setting under identical acquisition across groups. Aortic valve inflammation was evaluated in a blinded manner by assessing inflammatory cell infiltration and tissue remodeling on trichrome-stained sections.

### Plasmid vector cloning

Part of the 3’ untranslated region (UTR) of *MKI67* or *CBX5*, including wildtype (WT) or mutant seed sequence of miR-10b-5p, was purchased from Macrogen Co. (Seoul, South Korea) and was cloned into psiCHECK-2 vector (Promega, Madison, WI, USA), between the hRluc and poly(A) signal. Gibson Assembly was performed using NEBuilder HiFi DNA Assembly Master Mix (New England Biolabs, Ipswich, MA, USA) according to the manufacturer’s protocol. The WT or mutant seed sequences of miR-10b-5p are written in capital letters among the following DNA sequences.

*MKI67* 3’ UTR_WT:

actctgtaaagcatcatcatccttggagagactgagcactcagcaccttcagccacgatttcaggatcgcttccttgtgagccgctgcct ccgaaatctcctttgaagcccagacatctttctccagcttcagacttgtagatataactcgttcatcttcatttactttccactttgccccctgtc ctctctgtgttccccaaatcagagaatagcccgccatcccccaggtcacctgtctggattcctccccattcacccaccttgccaggtgca ggtgaggatggtgcaccagACAGGGTAgctgtcccccaaaatgtgccctgtgcgggcagtgccctgtctccacgtttgtttccc cagtgtctggcggggagccaggtgacatcataaatacttgctgaatgaatgcagaaatcagcggtactgacttgtactatattggctgcc atgatagggttctcacagcgtcatccatgatcgtaagggagaatgacattctgcttgagggagggaatagaaaggggcagggaggg gacatctgagggcttcacagggctgcaaagggtacagggattgcaccagggcagaacaggggagggtgtt

*MKI67* 3’ UTR_mutant:

actctgtaaagcatcatcatccttggagagactgagcactcagcaccttcagccacgatttcaggatcgcttccttgtgagccgctgcct ccgaaatctcctttgaagcccagacatctttctccagcttcagacttgtagatataactcgttcatcttcatttactttccactttgccccctgtc ctctctgtgttccccaaatcagagaatagcccgccatcccccaggtcacctgtctggattcctccccattcacccaccttgccaggtgca ggtgaggatggtgcaccagGCCTTCAAgctgtcccccaaaatgtgccctgtgcgggcagtgccctgtctccacgtttgtttccc cagtgtctggcggggagccaggtgacatcataaatacttgctgaatgaatgcagaaatcagcggtactgacttgtactatattggctgcc atgatagggttctcacagcgtcatccatgatcgtaagggagaatgacattctgcttgagggagggaatagaaaggggcagggaggg gacatctgagggcttcacagggctgcaaagggtacagggattgcaccagggcagaacaggggagggtgtt

*CBX5* 3’ UTR_WT:

gagatggaaaggatgttgcccatctgttaaaaagccaatagcaactgcctacctgttggggctttcccaaccttgttcagctctacccaggagaatACAGGGTttctggtgtggtcttctgggccaccctctgttgttcatcctcactcatcctcccagaattactccatcctttggaagatttgaaattttacactgaaatctgtcaagacactctttttagccccaggactttcggtattgcttttagcactccccacttctgcttctgtaaagtgatatcaatttgtgataaaatgacaagattattaactgtgcagatttgtagctatagtacatgaggatgatggggtagggtacaattgtctttattatcatatatggtatgtatgtatgatttctttccattcctcatatttagactgtatatttatgtaggtgtgagtgattgcctggtgcttgcttgtgccaaggtgctaggcaccctccaaccctgccaacttttgtggcctcccaaagcattcctgttaccaaagaggcttcaaacctgaccctcacttctcagtggacccgagtttcccttccatgccattattttcagtggggaagttttagaggtgagctgttggccacaatatcaattttaagtgttcatagcagttatgtctcctgcattcttggctcctggatttaccaccaagagtccccaaaatattaatgctcttccctttttctaccctcaaacttatagttgtatcttattttttaaaatgaattttcatggccaggcacagtggctaccgcctgtaatcccagcactttgggaggctgaggcaggagaattgcttgaagccagaagtttgagaccagcctgggcaatgtagtgagaccccctgtctct

*CBX5* 3’ UTR_mutant:

gagatggaaaggatgttgcccatctgttaaaaagccaatagcaactgcctacctgttggggctttcccaaccttgttcagctctacccaggagaatGTCTAAGttctggtgtggtcttctgggccaccctctgttgttcatcctcactcatcctcccagaattactccatcctttggaagatttgaaattttacactgaaatctgtcaagacactctttttagccccaggactttcggtattgcttttagcactccccacttctgcttctgtaaagtgatatcaatttgtgataaaatgacaagattattaactgtgcagatttgtagctatagtacatgaggatgatggggtagggtacaattgtctttattatcatatatggtatgtatgtatgatttctttccattcctcatatttagactgtatatttatgtaggtgtgagtgattgcctggtgcttgcttgtgccaaggtgctaggcaccctccaaccctgccaacttttgtggcctcccaaagcattcctgttaccaaagaggcttcaaacctgaccctcacttctcagtggacccgagtttcccttccatgccattattttcagtggggaagttttagaggtgagctgttggccacaatatcaattttaagtgttcatagcagttatgtctcctgcattcttggctcctggatttaccaccaagagtccccaaaatattaatgctcttccctttttctaccctcaaa cttatagttgtatcttattttttaaaatgaattttcatggccaggcacagtggctaccgcctgtaatcccagcactttgggaggctgaggcaggagaattgcttgaagccagaagtttgagaccagcctgggcaatgtagtgagaccccctgtctct

### Dual-luciferase reporter assay

HEK 293T cells were transfected with psiCHECK-2_MKI67 or CBX5 3’ UTR_WT or mutant using Lipofectamine 3000 (Invitrogen, Waltham, MA, USA) according to the manufacturer’s manual and treated with scramble or miR-10b-5p mimic the next day. A Dual-Luciferase Reporter Assay System (Promega, Madison, WI, USA) was used according to the manufacturer’s manual. Human Renilla luciferase signals were normalized by firefly luciferase signals.

### Enzyme-linked immunosorbent assay

Human IL-8/CXCL8 DuoSet ELISA (R&D Systems, Minneapolis, MN, USA) and DuoSet ELISA Ancillary Reagent Kit 2 (R&D Systems, Minneapolis, MN, USA) were used for CXCL8 ELISA. For the assay procedure, 100 μL of patient sera or standards were added per well and experimented according to the manufacturer’s protocol. Optical densities (ODs) were measured at 450 nm with a microplate reader (Agilent BioTek, Santa Clara, CA, USA), and ODs at 570 nm were subtracted to correct optical imperfections.

### Receiver operating characteristic (ROC) analysis

The diagnostic performance of serum CXCL8 for discriminating KD from febrile controls was evaluated using receiver operating characteristic (ROC) curve analysis. Sensitivity and specificity were computed across all possible thresholds, and an optimal cut-off value was determined by maximizing the sum of sensitivity and specificity. ROC analyses were performed using Prism 9 (San Diego, CA, USA).

### Chromatin immunoprecipitation

HCAECs or HEK293T cells were treated with an EGFP or HA-hCEBPA -expressing adenoviral vector (Ad-EGFP or Ad-HA-hCEBPA; VectorBuilder, Chicago, IL, USA) at a multiplicity of infection (MOI) of 5 or 1 overnight. Cells were briefly rinsed with PBS, and chromatin extraction and immunoprecipitation were performed using EZ-ChIP (17-371, Sigma-Aldrich, Burlington, MA, USA) and anti-HA antibody (2.5 μg; H9658, Sigma-Aldrich, Burlington, MA, USA), according to the manufacturer’s manual. After the final elution step, we performed qPCR on the immunoprecipitated DNA using the following primers. The amplicons were 152 and 172 bp each.

*CXCL8*_forward: TGCTTTCTTCTTCTGATAGACCAAACTC

*CXCL8*_reverse: TGTTAACAGAGTGAAGGGGCACA

*MMP10*_forward: ATGTAGAGAAAAGAATCCAAAGAGAGAAGC

*MMP10*_reverse:

TTAAAAGCAAAGAAGAGGAAGAGGGTAGGA

The cycle threshold (CT) values were normalized using the input chromatin values and Ad-EGFP-treatment values.

### *In vitro* binding assay

HEK293T cells were infected with Ad-HA-hCEBPA (VectorBuilder, Chicago, IL, USA) at an MOI of 1 overnight. Cells were briefly rinsed with PBS and lysed with 1% NP-40 buffer (50 mM Tris-HCl, pH 8.0; 150 mM NaCl; 1% NP-40; 1 mM EDTA). The extracted protein was quantified, and 200 μg of protein was used for the downstream experiments.

To biotinylate total DNA fragments of the *CXCL8* or *MMP10* promoter region, biotinylated primers flanking the fragments were produced, and PCR was performed. For the rest of the fragments, biotinylated double-stranded oligos were directly synthesized (BIONICS Co., Seoul, South Korea).

1 μg of DNA fragments were bound to 30 μL Dynabeads M-270 Streptavidin (65305, Invitrogen, Waltham, MA, USA) for 1 hour on a rotator at RT. The beads were washed 2 times with 1% NP-40 buffer. DNA-bound beads were bound to the 200 μg of protein for 2 hours on a rotator at RT. The beads were washed 3 times with 1% NP-buffer. The DNA-and-protein-bound beads were eluted with SDS sample buffer, and SDS-polyacrylamide gel electrophoresis (SDS-PAGE) was performed. We treated anti-HA antibody (1:10,000; H9658, Sigma-Aldrich, Burlington, MA, USA) as a primary antibody and peroxidase-conjugated Pierce goat anti-mouse antibody (1:20,000; 31430, Thermo Fisher Scientific, Waltham, MA, USA) as a secondary antibody. The signal was detected using a chemiluminescence system.

*CXCL8*_total:

TTACTCAGAAAGTTACTCCATAAATGTTTGTGGAACTGATTTCTATGTGAAGCACATGTGCCCCTTCACTCTGTTAACATGCATTAGAAAACTAAATCTTTTGAAAAGTTGTAGTATGCCCCCTAAGAGCAGTAACAGTTCCTAGAAACTCTCTAAAATGCTTAGAAAAAGATTTATTTTAAATTACCTCCCCAATAAAATGATTGGCTGGCTTATCTTCACCATCATGATAGCATCTGTAATTAACTGAAAAAAAATAATTATGCCATTAAAAGAAAATCATCCATGATCTTGTTCTAACACCTGCCACTCTAGTACTATA

*CXCL8*_fragment 1: TTACTCAGAAAGTTACTCCATAAATGTTTGTGGAACTGATTTCTATGTGA

*CXCL8*_fragment 2: GTGAAGCACATGTGCCCCTTCACTCTGTTAACATGCATTAGAAAACTAAA

*CXCL8*_fragment 3: AAATCTTTTGAAAAGTTGTAGTATGCCCCCTAAGAGCAGTAACAGTTCCT

*CXCL8*_fragment 4: TGTAATTAACTGAAAAAAAATAATTATGCCATTAAAAGAAAATCATCCAT

*MMP10*_total:

AAGAATTAAGTAGGTCAACGGAAATAAATTCCAAATATAGTCGTGGTCTAAATGTGGGAGTTGTTATTTACAAAGATATTAAAACAAATATTTCACTTCTTGAGTCCTTGTTTACAATAGCTATTATTGATTTTCAGATGGATTTACTGGCCGTGGAGTAGGGGTATTGTGAGAAGGACATGGAGCATGTAGAGAAAAGAATCCAAAGAGAGAAGCTTTGGTGTTCACTCATTCTCCATTTCCCAGTGTATAATTATGTTCAAAGTAACTGCTCCATAAAA CTTGTTAAATGGAAAAAATTAAATTGCATGTTAGACTGAAATGGTCCTCCTACCCTCTTCCTCTTCTTTGCTTTTAAAAATTCTTTTCCCCCTTTCTT

*MMP10*_fragment 1:

AGTAGGTCAACGGAAATAAATTCCAAATATAGTCGTGGTCTAAATGTGGG

*MMP10*_fragment 2:

TTGGTGTTCACTCATTCTCCATTTCCCAGTGTATAATTATGTTCAAAGTA

*MMP10*_fragment 3:

TATAATTATGTTCAAAGTAACTGCTCCATAAA

*MMP10*_fragment 4:

AAACTTGTTAAATGGAAAAAATTAAATTGCATGTTAGACTGAAATGGTCC

### Single-cell RNA-sequencing

The scRNA-seq experiment was performed by Macrogen Co. (Seoul, South Korea). Briefly, miRNA mimic-treated HCAECs were suspended as single cells and entered the Chromium Single Cell Gene Expression (10x Genomics) pipeline. Cells were individually barcoded in gel beads in emulsion (GEMs) and reverse transcribed. cDNAs were amplified with PCR. Gene expression libraries were constructed and 3’ sequenced (Illumina, San Diego, CA, USA). Sequencing results were provided as CellRanger output.

scRNA-seq data were analyzed using the Seurat package in R. The Seurat object was created with features of at least 3 cells and cells with at least 200 features. Cells with over 3000 and less than 6000 features and mitochondrial gene percent less than 5% were selected for quality control (QC). The data was log-normalized with a scale factor of 10,000. 2,000 highly variable features were selected using the variance-stabilizing transformation (VST) method. The data was then linearly transformed to the mean expression of 0 and the variance of 1. For the principal component analysis (PCA), 50 dimensions were selected. A shared nearest neighbor (SNN) method was used to cluster the cells with a resolution of 0.3.

Positive marker genes were found using the FindAllMarkers function in Seurat for the 5 clusters. Gene ontology (GO) analysis was performed using the enrichR package in R and Kyoto Encyclopedia of Genes and Genomes (KEGG) 2026 database, and each cluster was annotated as metabolic, inflammatory, protein-dynamic, others, and proliferative, respectively, according to the top 10 GO terms.

Trajectory analysis was performed using the monocle3 package in R. The Seurat object was converted to the CellDataSet object for monocle3, and the trajectory graph was learned using the learn_graph function in monocle3. The proliferative cluster was set as the starting cells. To find the gene expressions that change according to the pseudotime, we used the graph_test function.

### Bulk RNA sequencing

The bulk RNA sequencing was performed by Macrogen Co. (Seoul, South Korea). Briefly, 3 biological replicates of scramble or miRNA mimic-treated frozen HCAECs were lysed, and the whole RNAs were extracted. cDNAs were synthesized from the whole RNAs, and the sequencing library was constructed. All libraries were sequenced on the Illumina platform (Illumina, San Diego, CA, USA). The reads were filtered using FastQC and aligned to the reference sequence using HISAT2. Differentially expressed gene analysis was performed using DESeq2. Downstream gene set enrichment analysis was performed using the GSEA_4.4.0 program with human MSigDB hallmark gene sets with default parameters.

### Assay for transposase-accessible chromatin with sequencing

Bulk assay for transposase-accessible chromatin with sequencing (ATAC-seq) of HCAECs was performed by ebiogen Inc. (Seoul, South Korea) The construction of the library was performed using ATAC Seq Library Prep Kit for Illumina (Active Motif, CA, USA) according to the manufacturer’s instructions. Library quantity was checked using TapeStation HS D5000 Screen Tape (Agilent Technologies, CA, USA). Lastly, all libraries were sequenced on the NovaSeq 6000 with 100 bp paired-end reads (Illumina, CA, USA).

Low-quality DNA reads and adapter sequences were filtered out using fastp-0.23.1. Clean reads were aligned to the reference genome sequence using bowtie2-2.3.4.3. The mtDNA sequences and duplicated reads were removed using samtools-1.17 and picard-2.27.5, respectively. ENCODE blacklist regions were removed from the BAM file, and coordinates were shifted. Peaks in alignment files were identified using MACS2, and differential accessibility analysis was performed using the Python “conorm” package 1.2.0. Homer was used for motif analysis of peaks, and ChIPseeker was used for annotations.

### Statistics

Statistical analyses were performed using PASW Statistics 29 (IBM SPSS Statistics; IBM Corp., Chicago, IL, USA). Outliers were identified using Grubbs’ test, and values meeting the exclusion criterion were removed from the analysis. Normality was assessed using the Shapiro–Wilk test prior to hypothesis testing. Comparisons between two independent groups were performed using a two-tailed unpaired Student’s t-test for normally distributed data or the Mann–Whitney U test for nonparametric data. For analyses involving more than two groups, two-way ANOVA was applied. When a significant interaction was observed, group effects were further examined by stratified pairwise comparisons. Tukey’s HSD was used for post hoc. To evaluate changes in serum CXCL8 levels between the first visit (day 0) and the 8-week follow-up, Wilcoxon matched-pairs signed rank test was performed. A two-sided *p*-value < 0.05 was considered statistically significant.

## Results

### Circulating microRNA (miRNA) discovery identifies Kawasaki disease (KD)-associated candidates

Because rapid diagnosis is critical to prevent coronary artery aneurysm formation, a potentially life-threatening complication of KD, early identification is essential. However, there is still no established diagnostic marker that can reliably distinguish KD from other febrile illnesses in patients presenting to the ED or outpatient clinic with high fever.

Therefore, we enrolled patients who presented to the ED with high fever as the study population. Blood samples were obtained from patients who provided informed consent before a final diagnosis was established in the ED. Plasma and buffy coat fractions were separated and stored for subsequent analyses (Figure 1A). After completion of the diagnostic evaluation, patients were retrospectively classified into a febrile control group if a specific non-Kawasaki etiology was identified, or into the KD group if they were finally diagnosed with KD. To profile global miRNA changes at the time of ED presentation, we randomly selected five cases from the febrile control group and five from the KD group. Whole miRNA sequencing was performed on buffy coat specimens obtained from selected subjects in each study group, because the buffy coat is enriched in immune cell–derived molecules and is therefore expected to reflect the inflammatory milieu of KD. We identified 40 significantly upregulated miRNAs in KD samples (Figure 1B and 1C), suggesting broad activation of miRNA programs during acute KD.

**Figure 1.**
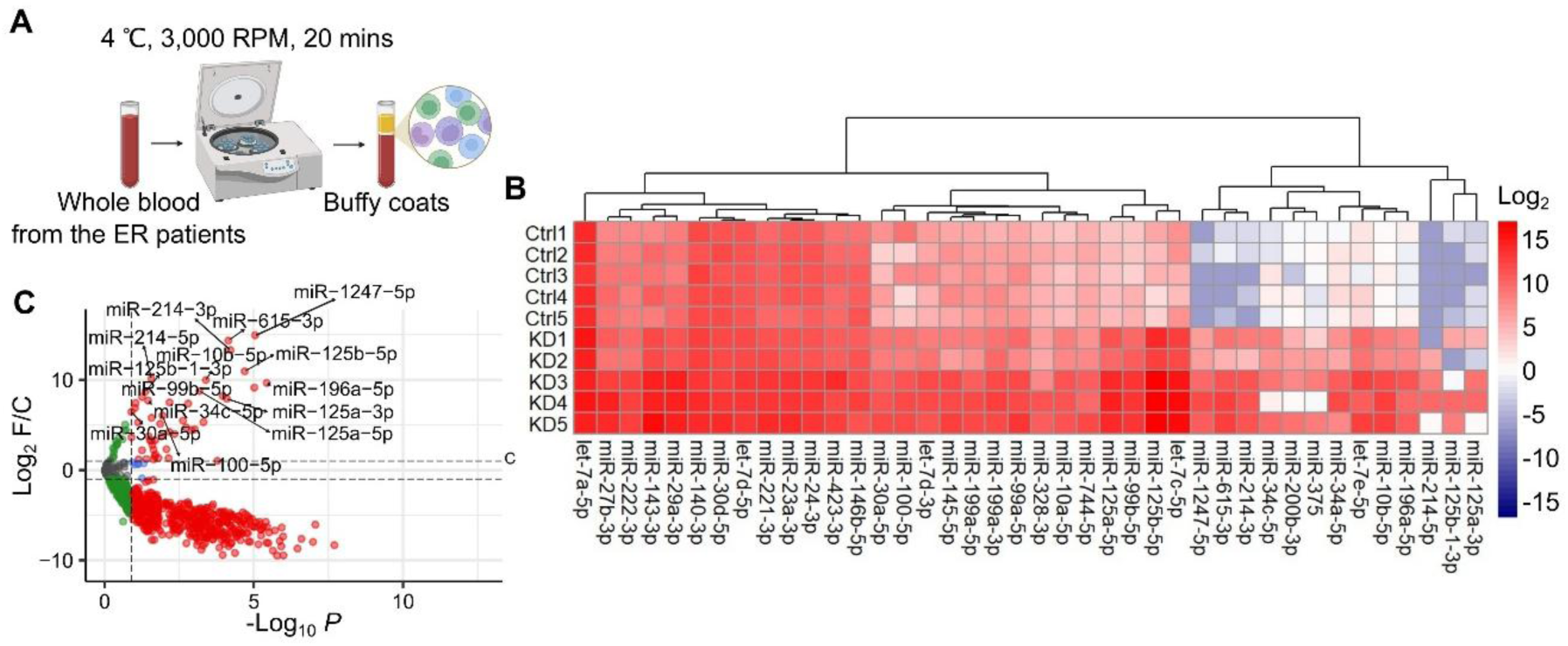
Circulating miRNA profiling identifies KD-associated candidates at emergency department presentation. **A**, Schematic of the study design. Whole blood was collected from patients presenting to the emergency department (ED) with acute febrile illness, centrifuged at 3,000 RPM for 20 minutes at 4°C, and buffy coat fractions were isolated for total RNA extraction and miRNA profiling. Patients were retrospectively classified into febrile control or KD groups based on the final diagnosis. **B**, Heatmap of significantly upregulated miRNAs in acute KD buffy coats. Log_2_ fold change was calculated using group means (KD vs. febrile controls) and miRNAs were hierarchically clustered. KD and febrile control samples are well separated, indicating a distinct miRNA signature during acute KD (n=5 per group). **C**, Volcano plot of differentially expressed miRNAs in buffy coats of KD patients versus febrile controls. Each point represents one detected miRNA. Fourteen candidate miRNAs selected for downstream validation—miR-10b-5p, miR-30a-5p, miR-34c-5p, miR-99b-5p, miR-100-5p, miR-125a-3p, miR-125a-5p, miR-125b-1-3p, miR-125b-5p, miR-196a-5p, miR-214-3p, miR-214-5p, miR-615-3p, and miR-1247-5p—are labelled.

We assumed that not all differentially expressed miRNAs would be equally functionally relevant. Specifically, among these upregulated miRNAs, we hypothesized that miRNAs with (i) consistent expression across KD patients (low inter-individual variance) and (ii) robust concentrations (high normalized counts) would have played an important role in KD progression. With these criteria, we filtered the upregulated miRNAs and finally decided to investigate further fourteen candidate miRNAs, which are miR-10b-5p, miR-30a-5p, miR-34c-5p, miR-99b-5p, miR-100-5p, miR-125a-3p, miR-125a-5p, miR-125b-1-3p, miR-125b-5p, miR-196a-5p, miR-214-3p, miR-214-5p, miR-615-3p, and miR-1247-5p. These miRNAs were selected because they combined upregulation with stable, high-abundance expression patterns, making them suitable candidates for the following validations.

### *In vivo* validation of miR-10b-5p as a pathogenic driver

To determine whether the fourteen miRNA candidates are functionally involved in the pathogenesis of KD, we generated two widely established murine models for KD-like vasculitis (LCWE and CAWS). Because both the LCWE and CAWS models recapitulate key features of human KD, including coronary arteritis and inflammatory cell infiltration in aortic roots^7,24^, it provides an *in vivo* platform to test whether candidate miRNAs are functionally required. Using this model, we aimed to test the *in vivo* reproducibility of the patient-derived miRNA signature and to narrow the candidate list to those most robustly linked to vasculitis.

We first intraperitoneally injected vehicle or LCWE into the BALB/c and collected whole blood samples at 4 weeks to evaluate changes in circulating miRNA levels (Figure 2A). At 4 weeks post-injection, we observed that several candidates identified in human KD were similarly increased in the LCWE model. Specifically, the relative blood expression levels of miR-10b-5p, miR-30a-5p, miR-99a-5p, miR-125a-3p, miR-196a-5p, miR-214-3p, and miR-615-3p were significantly higher in LCWE-treated mice compared with the vehicle-injected controls (Figure 2B, 2C, and S1). These data indicated that a subset of the patient-derived candidates is reproducibly induced in a KD-like vasculitis model, supporting their biological relevance and prioritizing them for further study.

**Figure 2.**
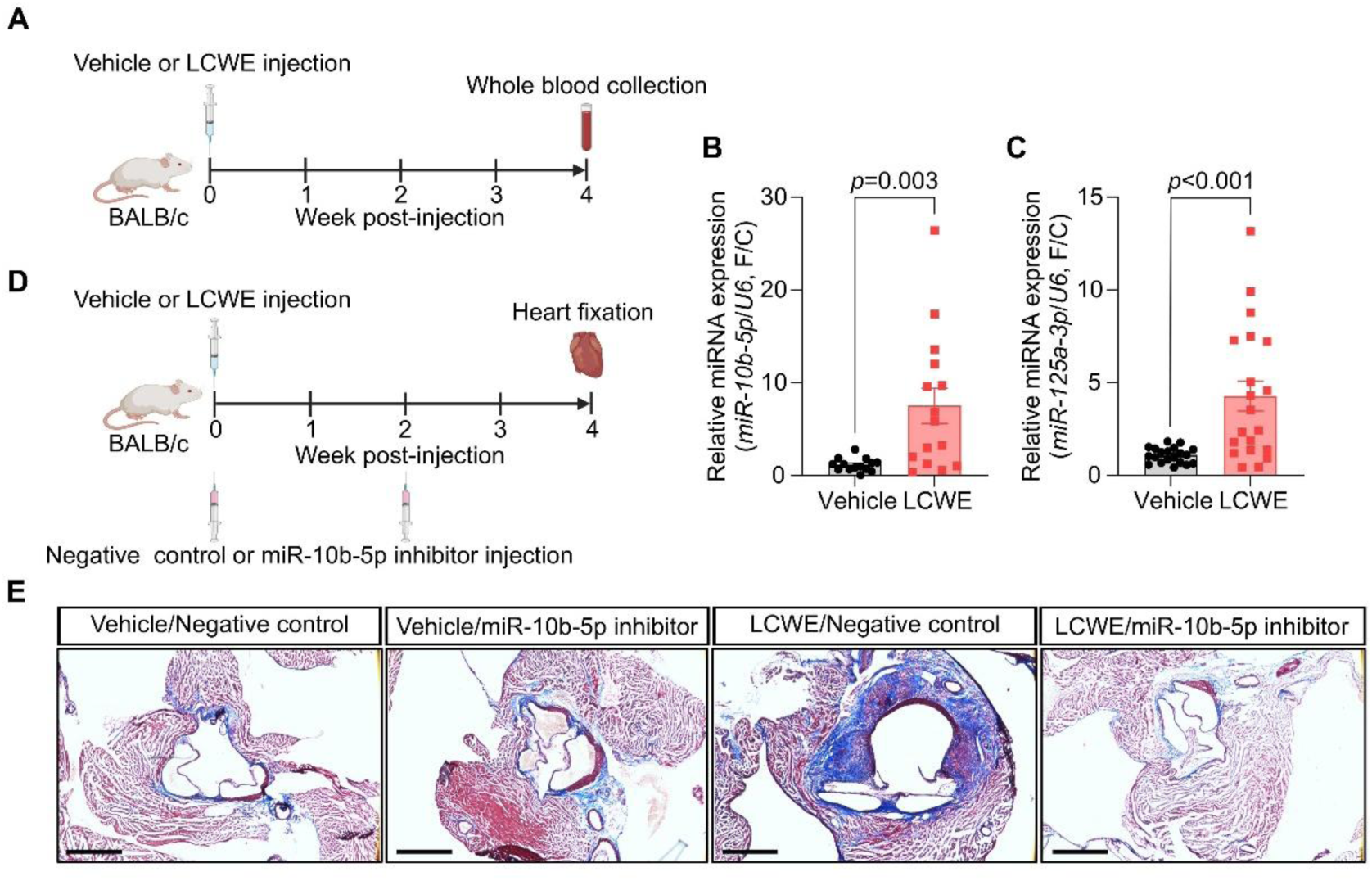
miR-10b-5p is required for KD-like vasculitis *in vivo*. **A**, Schematic of the LCWE vasculitis model. BALB/c mice received intraperitoneal injection of vehicle or LCWE and whole blood was collected at 4 weeks post-injection for circulating miRNA quantification. **B** and **C**, Relative blood expression levels of miR-10b-5p (**B**) and miR-125a-3p (**C**) in vehicle- or LCWE-injected mice at 4 weeks post-injection, measured by reverse transcription and quantitative polymerase chain reaction (RT-qPCR). LCWE challenge significantly elevated both miR-10b-5p and miR-125a-3p (Mann–Whitney U test; **B**: vehicle, n=14; LCWE, n=15; **C**: vehicle, n=20; LCWE, n=20). **D**, Schematic of the loss-of-function experiment. Mice undergoing LCWE modeling were co-administered a locked nucleic acid (LNA)-based antagomir targeting miR-10b-5p (LNA-anti-miR-10b-5p) or a matched negative control via tail-vein injection, with a second dose at week 2. Hearts were fixed at 4 weeks post-injection for histological assessment of aortic root inflammation. **E**, Representative Masson’s trichrome staining (MTS) images of aortic valve sections from each experimental group. LNA-anti-miR-10b-5p attenuated LCWE-induced inflammatory cell infiltration and tissue remodeling at the aortic root. Scale bars, 500 μm.

To strengthen the *in vivo* validation and to reduce the possibility that the observed miRNA changes were specific to the LCWE stimulus, we additionally tested the candidates using another *in vivo* model system: the *Candida albicans* water-soluble fraction (CAWS) model. CAWS is a polysaccharide-rich extract from *Candida albicans* that, when administered intraperitoneally, induces coronary arteritis and vascular inflammation similar to human KD^24^. Because these two models trigger vasculitis through distinct inflammatory cues, miRNAs induced in both systems are more likely to represent core disease-related regulators rather than model-specific byproducts. We injected vehicle or CAWS into the BALB/c and collected whole blood samples at 4 weeks post-injection (Figure S2). At this time point, blood levels of miR-10b-5p, miR-30a-5p, and miR-125a-3p were significantly elevated compared with the vehicle-injected group (Figure S2A, S2B, and S2D). This further supported the reproducibility of miRNA induction during KD-like vasculitis.

Among the three validated miRNAs, we recalled the changes in KD patients. Notably, miR-30a-5p (5.30) showed a minimal log_2_ fold change in the human sequencing dataset than miR-10b-5p (9.97) and miR-125a-3p (7.70), suggesting that miR-10b-5p and miR-125a-3p may be more strongly associated with KD in patients and more influential in disease pathogenesis.

Based on this cross-species consistency and the magnitude of differential expression, we hypothesized that miR-10b-5p or miR-125a-3p might play a functional role in promoting vascular inflammation.

To directly test whether miR-10b-5p or miR-125a-3p is required for KD-like vasculitis *in vivo*, we performed loss-of-function experiments in the LCWE model using a locked nucleic acid (LNA)-based inhibitory oligonucleotide (Figure 2D). We intravenously injected a negative control, LNA-anti-miR-10b-5p or LNA-anti-miR-125a-3p, to suppress miRNA respectively during LCWE-induced vascular inflammation. Surprisingly, inhibition of miR-10b-5p led to a reduction in aortic root inflammation compared with control animals, whereas LNA-anti-miR-125a-3p failed to mitigate inflammation progression, which indicates that miR-10b-5p contributes functionally to the development of KD-like vasculitis *in vivo* (Figure 2E). These *in vivo* validation and inhibition experiments support a model in which miR-10b-5p is consistently induced across KD-like settings and is required for full progression of aortic root inflammation in the LCWE model.

### miR-10b-5p drives inflammatory reprogramming of the coronary endothelium

We next sought to define the vascular cell program altered by miR-10b-5p and determine how this miRNA could mechanistically promote KD-associated vascular inflammation. Among the numerous cellular components of the vasculature, endothelial cells-which form the inner lining of the vessel wall and directly encounter the circulating inflammatory mediators, have known as key initiators and amplifiers of immune activation in KD vasculitis^13^. Given that miR-10b-5p is elevated in the circulation of KD patients and is functionally required for vasculitis *in vivo*, we asked whether increased miR-10b-5p is sufficient to reprogram coronary artery endothelial cells toward a disease-prone state.

To test this, we established a cellular model of miR-10b-5p-treated human primary coronary artery endothelial cells (HCAECs). To capture cell-state heterogeneity and identify transcriptional programs altered at the single-cell level, we performed single-cell RNA sequencing (scRNA-seq). Dimensionality reduction and visualization using uniform manifold approximation and projection (UMAP) revealed that scramble or miRNA-treated HCAECs were apparently separated into distinct distributions, indicating a substantial miR-10b-5p-driven shift in transcriptional identity (Figure 3A and 3B). We annotated clusters according to the gene ontology (GO) analysis results, which were metabolic, inflammatory, protein-dynamic, others, and proliferative, respectively (Figure 3B and S3).

**Figure 3.**
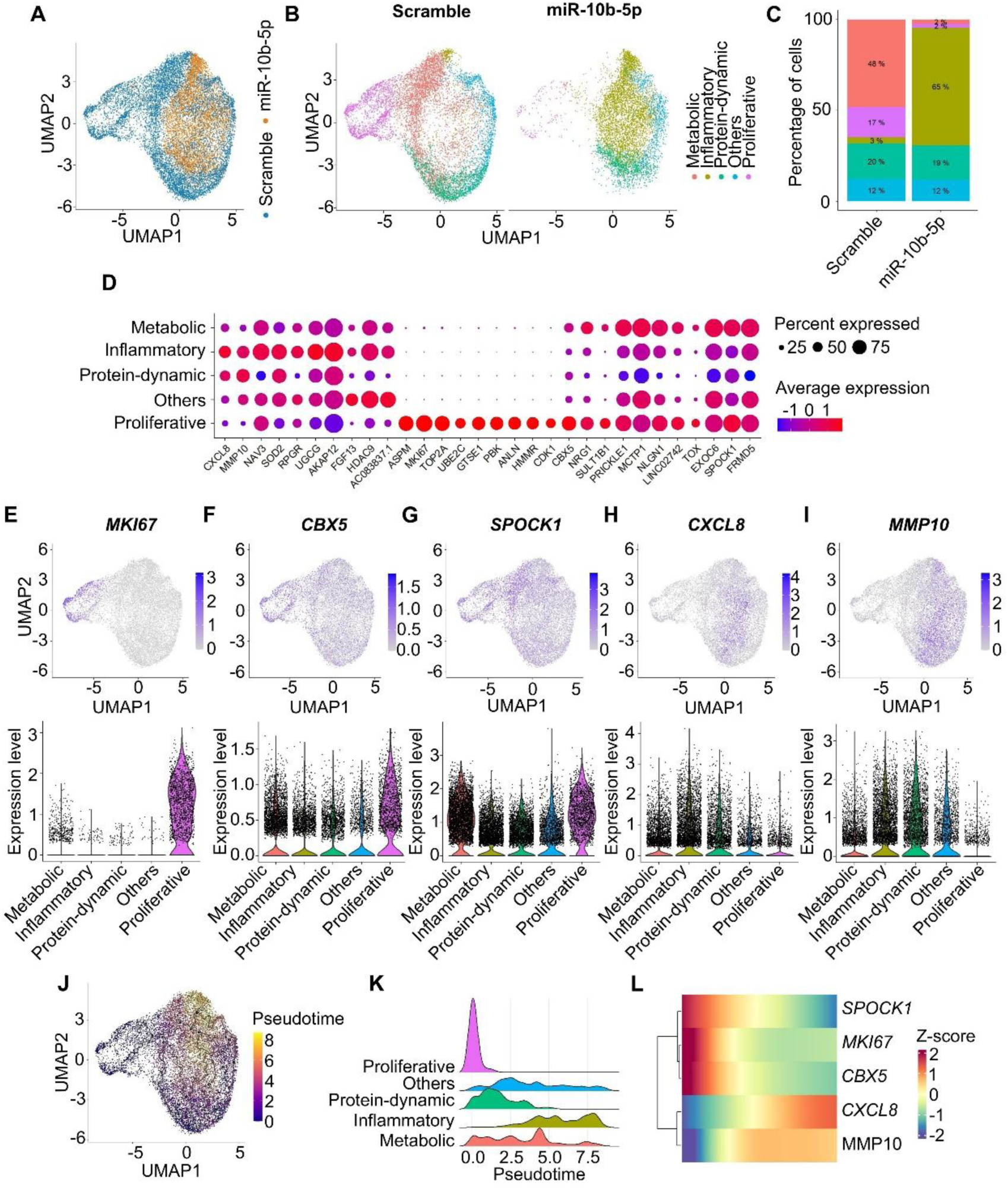
miR-10b-5p reprograms human coronary artery endothelial cells toward a pro-inflammatory state. **A**, UMAP visualization of HCAECs colored by treatment condition (scramble or miR-10b-5p mimic). miR-10b-5p treatment shifts the global transcriptomic distribution, indicating a substantial cell-state transition. **B**, UMAP plots colored by annotated cluster identity (metabolic, inflammatory, protein-dynamic, others, and proliferative), split by treatment condition to highlight miR-10b-5p-dependent cluster redistribution. **C**, Bar plot showing the percentage of cells in each cluster per condition. miR-10b-5p markedly expanded the inflammatory cluster (3% to 65%) while depleting the metabolic (48% to 2%) and proliferative (17% to 2%) clusters. **D**, Dot plot showing expression patterns of representative marker genes across clusters. Dot size indicates the percentage of expressing cells and color intensity indicates average expression level. **E**–**I**, Feature plots (upper row) and violin plots (lower row) showing cluster-restricted expression of *MKI67* (**E**), *CBX5* (**F**), *SPOCK1* (**G**), *CXCL8* (**H**), and *MMP10* (**I**). *MKI67* is predominantly expressed in the proliferative cluster; *CBX5* and *SPOCK1* are enriched in proliferative and metabolic clusters; *CXCL8* and *MMP10* are selectively expressed in the inflammatory cluster. **J**, UMAP plot colored by pseudotime score from trajectory analysis. The inferred trajectory originates in the proliferative cluster and progresses through the metabolic and protein-dynamic clusters toward the inflammatory cluster. **K**, Ridge plot showing the pseudotemporal distribution of each cluster, consistent with a directional transition from proliferative/metabolic toward inflammatory states. **L**, Heatmap of Z-scored expression of *SPOCK1*, *MKI67*, *CBX5*, *CXCL8*, and *MMP10* along pseudotime. Genes were hierarchically clustered, illustrating coordinated downregulation of proliferative/metabolic genes and upregulation of inflammatory genes during the transition.

miR-10b-5p treatment markedly altered the relative composition of these cell states. The miR-10b-5p group displayed a lower percentage of cells in the metabolic (2 vs. 48%) and proliferative clusters (2 vs. 17%), and a higher percentage of cells in the inflammatory cluster (65 vs. 3%; Figure 3C) whereas both protein-dynamic and other clusters were minimally changed. Among the three prominently altered clusters, the metabolic and proliferative clusters were enriched in scramble-treated HCAECs, whereas an inflammatory cluster emerged specifically upon miR-10b-5p treatment. This shift suggested that miR-10b-5p primarily targets genes characteristic of the proliferative and metabolic programs (Figure 3D). We therefore screened the 3′ untranslated regions (3′ UTRs) of cluster-defining genes for the miR-10b-5p seed sequence (ACAGGGTA) and identified *MKI67* (proliferative cluster), *SPOCK1* (shared by proliferative and metabolic clusters), and *CBX5* (shared by proliferative and metabolic clusters) as *in silico* candidates. In contrast, KD-associated inflammatory drivers were selectively expressed in the inflammatory cluster; thus, we prioritized *CXCL8* and *MMP10* as potential mediators of KD progression. Feature plots showed cluster-restricted expression patterns: *MKI67* was largely confined to the proliferative cluster (Figure 3E), *CBX5* was expressed across both proliferative and metabolic clusters with relatively broader distribution (Figure 3F), *SPOCK1* was enriched in the metabolic cluster (Figure 3G), and the chemotactic factors *CXCL8* and *MMP10* were abundant in the inflammatory cluster (Figure 3H and 3I). Indeed, when the feature plot was stratified into scramble and miR-treated groups, *MKI67*, *CBX5*, and *SPOCK1* are abundant under the scramble condition, whereas *CXCL8* and *MMP10* are enriched in the miR-treated group (Figure S4).

We hypothesized that miR-10b-5p drives a transition of proliferative and metabolic endothelial states toward an inflammatory cluster by directly targeting key genes. To further confirm that these dynamic changes were from a coordinated transition rather than static cluster differences, we performed downstream trajectory analysis. According to the inferred pseudotemporal axis, HCAECs transitioned from the proliferative cluster to the metabolic cluster under miR-10b-5p treatment and ultimately the inflammatory cluster (Figure 3J). The gene expression dynamics further demonstrated the dynamic transition accordingly (Figure 3K and 3L). These results indicate that miR-10b-5p is sufficient to reprogram human coronary artery endothelium at the transcriptome level, shifting cells away from homeostatic proliferative/metabolic programs and toward a pro-inflammatory phenotype.

### miR-10b-5p reprograms coronary vascular wall cells toward an inflammatory state

Although scRNA-seq is highly sensitive for resolving cell-to-cell heterogeneity, it is generated from a relatively limited number of cells and may be less powered to detect subtle, low-amplitude expression changes^25^. We therefore performed bulk RNA sequencing (RNA-seq) to complement the single-cell analyses and capture global transcriptomic shifts averaged across a large cell population. Because KD vasculitis involves multiple vascular wall cell types and vascular smooth muscle cells are also important contributor^7,26,27^, we analyzed both HCAECs (Figure 4A through 4F) and primary human coronary artery smooth muscle cells (HCASMCs) (Figure 4G through 4L) following miR-10b-5p treatment. Consistent with the scRNA-seq findings, the metabolic/proliferative to inflammatory transition was also recapitulated in the bulk RNA-seq of both HCAECs and HCASMCs (Figure 4A through 4L). In gene set enrichment analysis (GSEA), miRNA-treated HCAECs were enriched in epithelial mesenchymal transition (EMT, NES = 1.71, FDR = 0.014) inflammatory response (NES = 1.69, FDR = 0.015) and protein secretion (NES = 1.51, FDR = 0.033), and were less enriched in fatty acid metabolism (NES = −1.35, FDR = 0.084) and cell cycle (NES = −2.74, FDR = 0.000) (Figure 4B through 4F and S5A). These findings suggested that miR-10b-5p induced cell-cycle arrest in HCAECs and promoted differentiation into secretory phenotype associated with inflammatory responses. Similar enrichment patterns were further observed in HCASMCs (Figure 4H through 4L and S5B). miR-10b-5p-treated HCASMCs exhibited marked pathway-level alterations, with significant enrichment of epithelial–mesenchymal transition (NES = 2.26, FDR = 0.000) and TNFα signaling via NF-κB (NES = 1.54, FDR = 0.027), alongside significant depletion of the G2/M checkpoint (NES = −2.46, FDR = 0.000) and mitotic spindle (NES = −1.91, FDR = 0.001). GO analysis of negatively regulated DEGs further indicated that miR-10b-5p robustly suppresses pathways involved in cell-cycle progression and chromatin organization, predominantly in HCAECs (Figure S6), supporting a shared miR-10b-5p-driven transcriptional reprogramming across the vascular wall.

**Figure 4.**
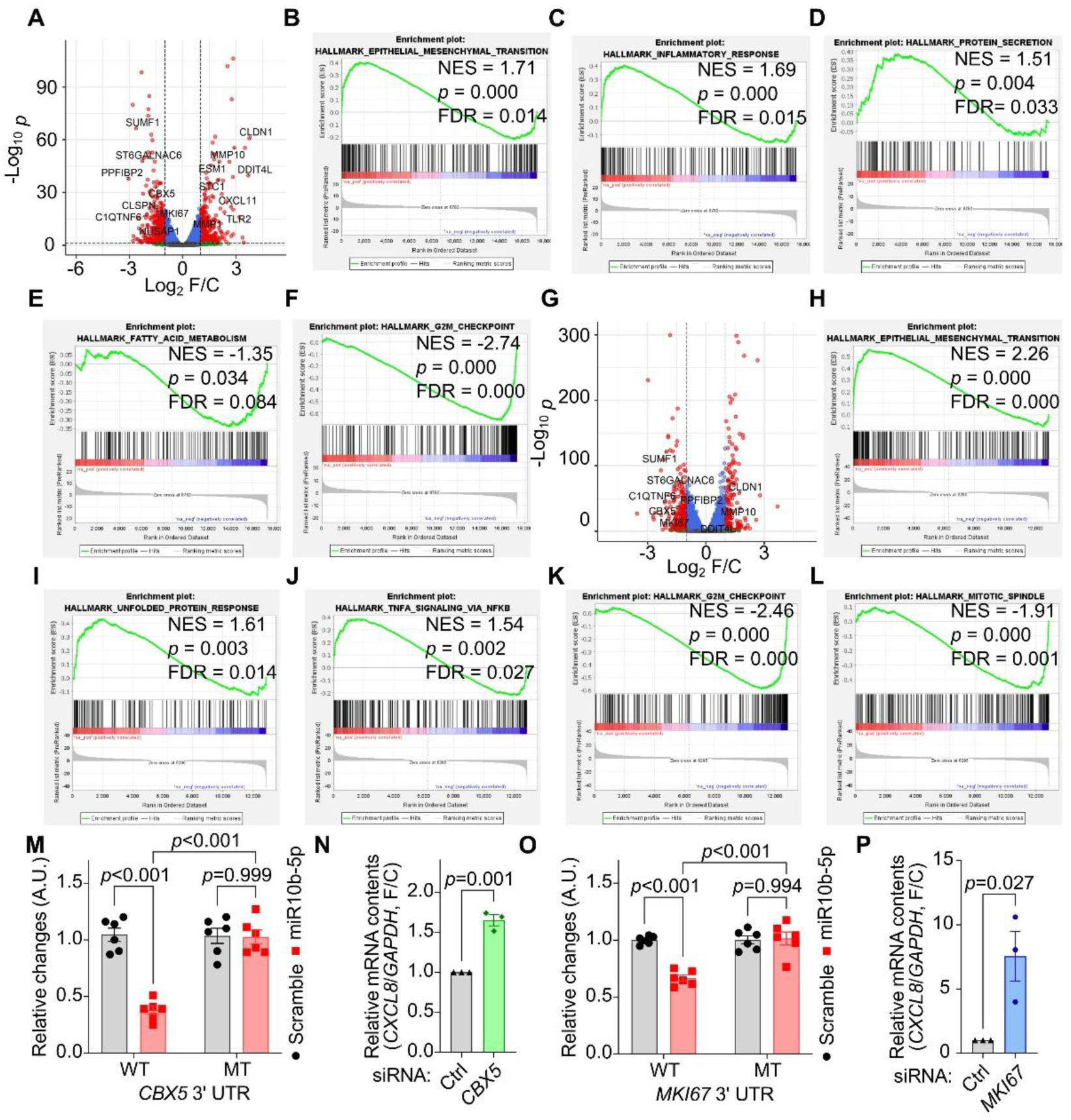
miR-10b-5p drives transcriptional reprogramming in coronary endothelial and smooth muscle cells and directly targets *CBX5* and *MKI67*. **A** and **G**, Volcano plots showing differentially expressed genes (DEGs) from bulk RNA-seq of miR-10b-5p-treated HCAECs (**A**) and HCASMCs (**G**). *MKI67* and *CBX5* are downregulated in both cell types, indicating suppression of proliferative programs. Red and blue dots denote significantly upregulated and downregulated genes, respectively. **B**–**F**, Gene set enrichment analysis (GSEA) plots from miR-10b-5p- versus scramble-treated HCAECs. miR-10b-5p enriched epithelial–mesenchymal transition (NES=1.71, FDR=0.014) (**B**), inflammatory response (NES=1.69, FDR=0.015) (**C**), and protein secretion (NES=1.51, FDR=0.033) (**D**) pathways, while depleting fatty acid metabolism (NES=−1.35, FDR=0.084) (**E**) and G2/M checkpoint (NES=−2.74, FDR<0.001) (**F**) pathways. **H**–**L**, GSEA plots from miR-10b-5p-treated HCASMCs, showing enrichment of epithelial–mesenchymal transition (NES=2.26, FDR<0.001) (**H**), unfolded protein response (NES=1.61, FDR=0.014) (**I**), and TNFα signaling via NF-κB (NES=1.54, FDR=0.027) (**J**), and depletion of G2/M checkpoint (NES=−2.46, FDR<0.001) (**K**) and mitotic spindle (NES=−1.91, FDR=0.001) (**L**) pathways. NES, normalized enrichment score; FDR, false discovery rate. **M** and **O**, Dual-luciferase reporter assay using wildtype (WT) or seed-sequence-mutant (MT) 3′ UTR constructs of *CBX5* (**M**) and *MKI67* (**O**) under scramble or miR-10b-5p mimic treatment. miR-10b-5p suppressed luciferase activity from WT but not MT reporters, confirming direct miRNA–target interactions (two-way ANOVA with Tukey’s HSD post hoc test; n=6 per group). **N** and **P**, Relative *CXCL8* mRNA levels in HCAECs following siRNA-mediated knockdown of *CBX5* (**N**) or *MKI67* (**P**). Knockdown of either target upregulates *CXCL8* expression, recapitulating the pro-inflammatory effect of miR-10b-5p (Student’s t-test; n=3 per group).

We next asked whether miR-10b-5p directly suppresses genes that maintain proliferative or metabolic states in vascular cells, thereby providing a mechanistic entry point into the observed global state transition. We found that *CBX5* and *MKI67* are strong candidate direct targets (Figure 3), each containing a miR-10b-5p seed sequence within their 3’ UTRs. Using a dual-luciferase reporter assay, we observed a significant reduction in luciferase activity when the 3′ UTR contained the predicted miR-10b-5p seed sequences. In contrast, miR-10b-5p failed to suppress luciferase activity when the seed sequences were replaced with random motifs. These findings confirmed that *CBX5* and *MKI67* are directly targeted by miR-10b-5p (Figure 4M and 4O), establishing that these genes are *bona fide* miR-10b-5p targets.

Next, we investigated how repression of these direct targets connects to the emergence of an inflammatory endothelial program. Notably, we found that *CXCL8*, encoding a neutrophil-attracting chemokine, is indirectly upregulated when *CBX5* and *MKI67* were suppressed (Figure 4N and 4P). This suggests that miR-10b-5p can promote a pro-inflammatory chemokine output not by directly targeting *CXCL8*, but by repressing upstream regulators associated with proliferative/chromatin programs, thereby tipping the transcriptional balance toward inflammatory activation.

These results support a model in which miR-10b-5p directly downregulates its target genes including *CBX5* and *MKI67*, drives a transition from proliferative/metabolic states to an inflammatory state in coronary endothelial and smooth muscle cells, and ultimately increases inflammatory chemokine expression such as *CXCL8*.

### CEBPA is the key transcriptional mediator of miR-10b-5p-induced *CXCL8* activation

We performed GSEA and GO enrichment analysis using transcriptomic changes induced by miR-10b-5p treatment both in HCAECs and HCASMCs (Figure 4, S5 and S6). The results indicated that miR-10b-5p induced cell-cycle arrest and disrupted chromatin remodeling, which are predominantly observed in HCAECs. To determine whether miR-10b-5p-induced transcriptional reprogramming was accompanied by chromatin remodeling, we next profiled chromatin accessibility in miR-10b-5p-treated HCAECs using assay for transposase-accessible chromatin using sequencing (ATAC-seq). Globally, the open chromatin landscapes appeared similar in scramble or miRNA-treated HCAECs with only subtle differences in overall accessibility patterns (Figure 5A). However, motif enrichment analysis using HOMER revealed that regions exhibiting newly increased chromatin accessibility upon miRNA treatment were most strongly enriched for CTCF and BORIS (also known as CTCFL) binding motifs (Figure 5B and 5C). GO analysis for the upregulated differentially accessible regions (DARs) displayed hormone receptors and metabolism-related terms (Figure 5D). These suggest that the miRNA may drive higher-order, global changes in chromatin architecture (i.e, enhancer-promoter loop) that extend beyond the local open-chromatin landscape detected in two dimensions, ATAC-seq.

**Figure 5.**
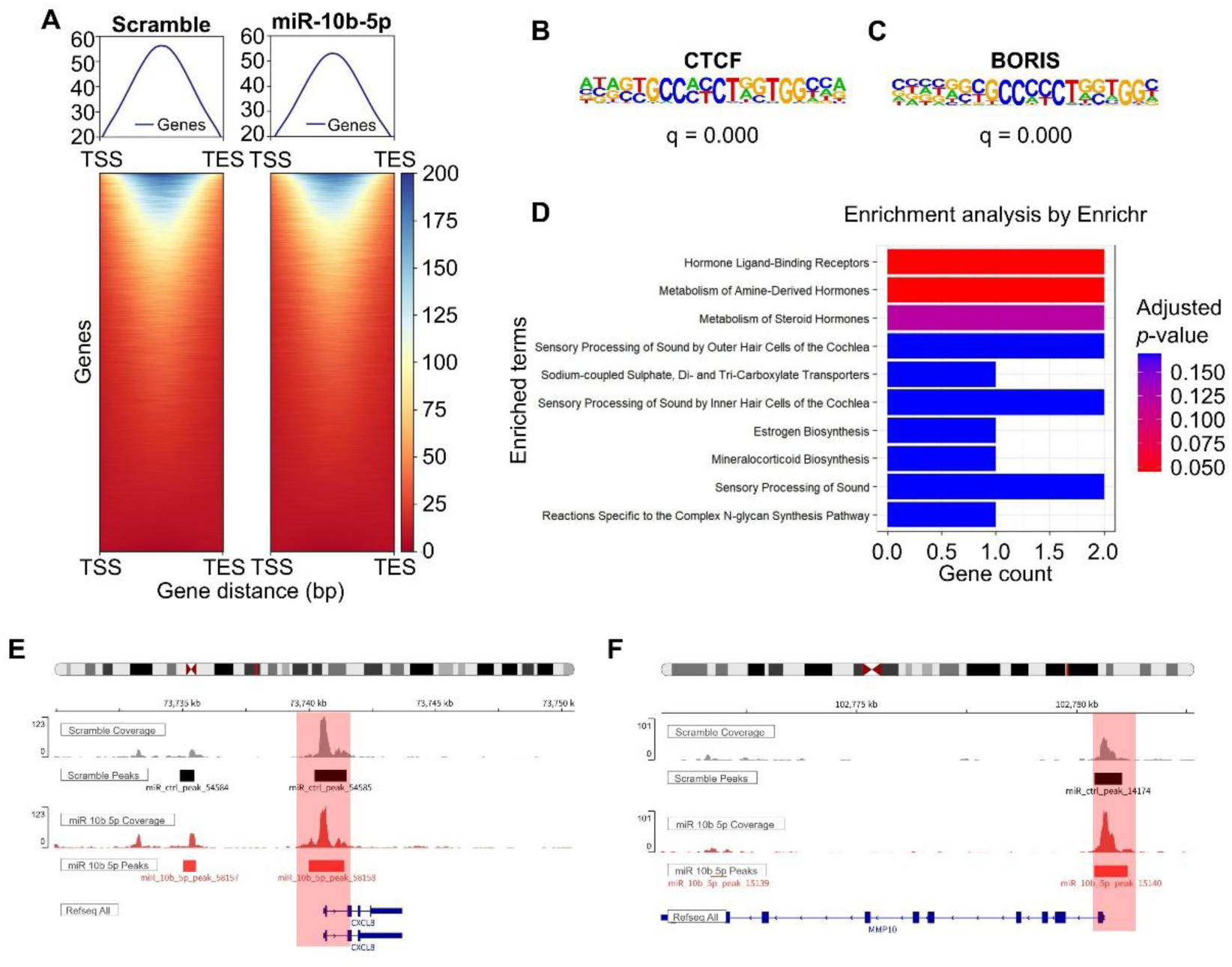
miR-10b-5p remodels chromatin accessibility in HCAECs. **A**, Genome-wide chromatin accessibility profiles of scramble- and miR-10b-5p mimic-treated HCAECs, displayed as average signal across all genes from transcription start site (TSS) to transcription end site (TES). Overall open chromatin is modestly reduced under miR-10b-5p treatment. **B** and **C**, Top enriched transcription factor binding motifs identified by HOMER in differentially accessible regions (DARs) of miR-10b-5p-treated HCAECs: CTCF (**B**; q=0.000, target 16.19%, background 9.24%) and BORIS (CTCFL) (**C**; q=0.000, target 16.23%, background 9.72%). **D**, Gene ontology (GO) analysis results of upregulated DARs in miR-10b-5p-treated HCAECs compared with scramble. Hormone ligand-binding receptors and metabolism of amine-derived hormones were enriched. **E** and **F**, Integrative Genomics Viewer (IGV) tracks showing chromatin accessibility at the *CXCL8* (**E**) and *MMP10* (**F**) loci in scramble- and miR-10b-5p-treated HCAECs. Shaded regions indicate DARs; accessibility is increased under miR-10b-5p.

Chromatin accessibility was not uniformly reduced by miRNA mimic treatment. More specifically, accessibility at cell cycle-associated loci was reduced under miR-10b-5p treatment. Representative examples included decreased accessibility around canonical proliferation-related genes such as *PCNA* and *CDK1* (Figure S7), matching the depletion of proliferative transcriptional programs observed in scRNA-seq and bulk RNA-seq. In contrast, loci associated with inflammatory effectors became more accessible, including increased chromatin accessibility at inflammatory genes such as *CXCL8* and *MMP10* (Figure 5E and 5F). Prominent peaks were detected in putative promoter regions of *CXCL8* and *MMP10*, along with broadly increased chromatin accessibility across their gene bodies (Figure 5E and 5F). These changes—closure of cell-cycle regulatory regions and opening of inflammatory regulatory regions—supported the conclusion that miR-10b-5p drives a pro-inflammatory shift in coronary endothelium not only at the transcriptomic level but also at the chromatin accessibility, implying durable remodeling of gene regulatory architecture.

We then sought to identify transcription factors that could bind newly accessible chromatin regions and activate the inflammatory gene program in the presence of miR-10b-5p, focusing on *CXCL8* and *MMP10* as a downstream effector identified in our transcriptomic analyses. Using motif analysis with the JASPAR database, we discovered putative binding sites for several transcription factors. Glucocorticoid receptors, CCAAT/enhancer-binding protein, Transcription Factor II D, GATA-binding protein, etc. To validate whether the candidate transcription factors were *bona fide* regulators of inflammatory gene expression, we co-transfected miR-10b-5p with siRNAs targeting each candidate transcription factor individually and quantified *CXCL8* and *MMP10* mRNA levels by qPCR. Interestingly, significant reductions in *CXCL8* and *MMP10* expression were observed only upon CEBPA (also known as C/EBPα) knockdown (Figure 6G, 6H). Although CEBP family subtypes share highly similar DNA-binding motifs, only CEBPA, unlike CEBPB or CEBPD, exerted a significant effect on *CXCL8* and *MMP10* expression (Figure S8).

**Figure 6.**
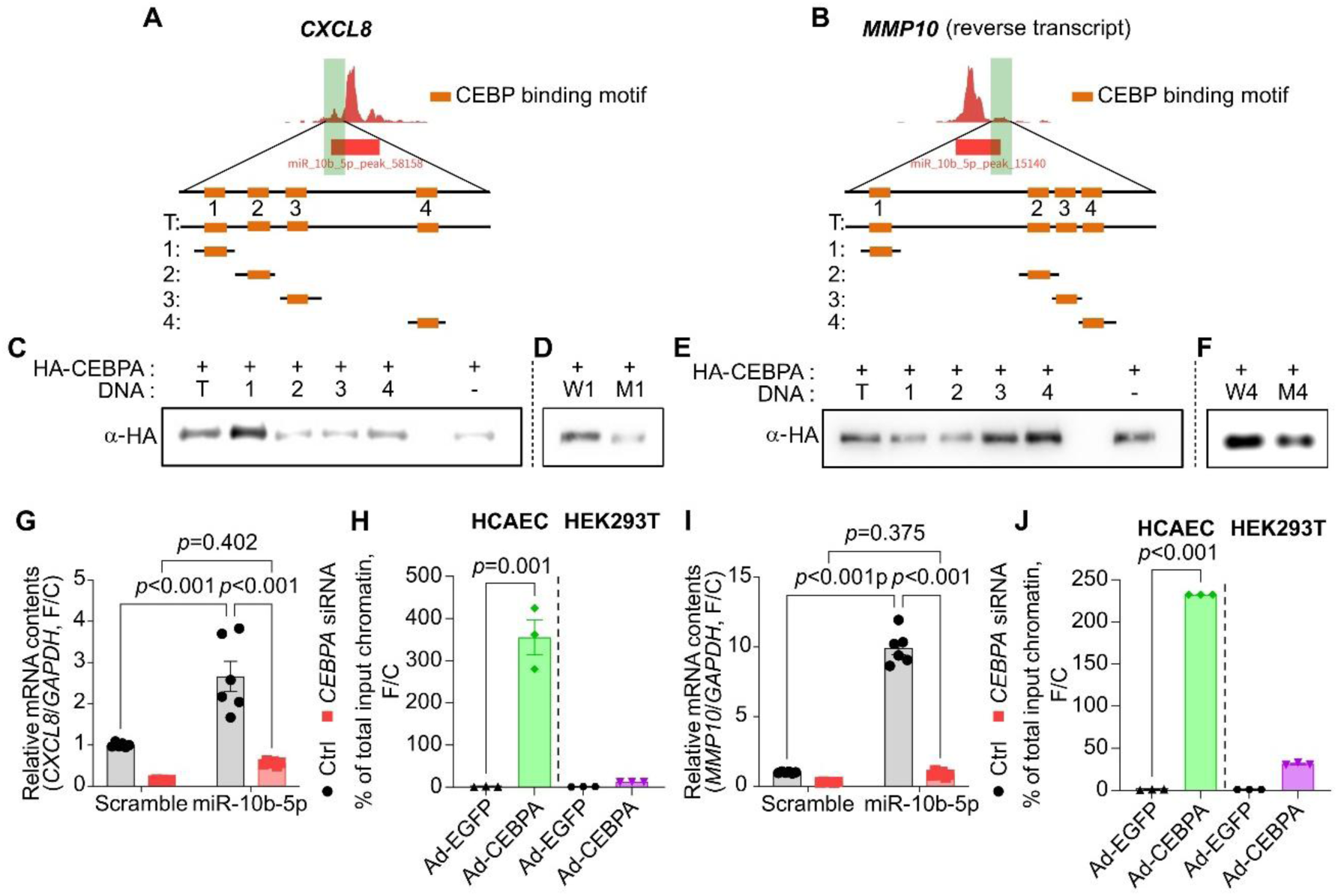
miR-10b-5p activates the CEBPA–*CXCL8*/*MMP10* transcriptional program. **A** and **B**, *CXCL8* (**A**) and *MMP10* (**B**) DARs in miR-10b-5p-treated HCAECs are shaded in red. These regions contained four CEBPA binding motif sites according to the JASPAR database. We synthesized and biotinylated the whole DAR fragments (about 300 bp each) containing all four binding motifs and the four DNA fragments (about 50 bp each) containing each binding motif, and used them for the in vitro binding assays. **C** and **E**, *In vitro* binding assay results for CEBPA binding to the *CXCL8* (**C**) and *MMP10* (**E**) promoter regions. CEBPA predominantly binds motif 1 of the *CXCL8* promoter, and motifs 3 and 4 of the *MMP10* promoter. **D** and **F**, *In vitro* binding assay confirming binding specificity: substitution of the CEBPA motif with a random sequence (M1/M4) abolishes binding compared with the wildtype (W1/W4) fragment in the *CXCL8* (**D**) and *MMP10* (**F**) promoters. **G** and **I**, Relative *CXCL8* (**G**) and *MMP10* (**I**) mRNA levels in HCAECs transfected with control siRNA (siCtrl) or *CEBPA* siRNA under scramble or miR-10b-5p conditions. *CEBPA* knockdown abolishes miR-10b-5p-induced upregulation of both genes (two-way ANOVA with Tukey’s HSD post hoc test; n=6 per group). **H** and **J**, CEBPA ChIP-qPCR at the *CXCL8* (**H**) and *MMP10* (**J**) promoters in HCAECs and HEK293T cells overexpressing Ad-HA-hCEBPA or Ad-EGFP. CEBPA occupancy is substantially higher in HCAECs than in HEK293T cells, supporting cell-type-specific engagement at inflammatory loci (Student’s t-test; n=3 per group).

We next investigated whether CEBPA directly binds to the newly accessible chromatin regions within the *CXCL8* and *MMP10* loci. To further validate direct binding, we performed an *in vitro* binding assay using CEBPA protein and target DNA sequences derived from *CXCL8* or *MMP10* promoter regions. As a result, CEBPA successfully bound to the *CXCL8* promoter (Figure 6C and 6D) or *MMP10* promoter (Figure 6E and 6F). Because each promoter contains four putative binding sites (Figure 6A and 6B), we repeated the assay using shorter DNA fragments flanking a single motif to validate the specific loci. We found that CEBPA predominantly binds the first motif within the *CXCL8* promoter, whereas it binds primarily to the third and fourth motifs in the *MMP10* promoter (Figure 6C and 6E). Of course, substitution of the binding motif within each short fragment with a random sequence abolished CEBPA binding (Figure 6D and 6F).

Chromatin immunoprecipitation (ChIP) further confirmed that CEBPA bound to the *CXCL8* promoter region in the cells (Figure 6H and 6J), especially in the HCAEC contexts. These findings demonstrate that miR-10b-5p induces chromatin remodeling and that CEBPA binds to newly accessible chromatin regions to drive expression of inflammatory genes such as *CXCL8* and *MMP10* in endothelial reprogramming.

### Serum CXCL8 is elevated in acute KD and shows diagnostic utility at first presentation

We have shown that miR-10b-5p is significantly elevated in samples from patients presenting to the ED with KD compared with other febrile illnesses. We further demonstrated that miR-10b-5p induces cell-cycle arrest and modulates chromatin remodeling in HCASMCs and HCAECs, with particularly pronounced effects in HCAECs. We also found that CEBPA acts as a key transcription factor binding to the newly accessible chromatin regions, thereby promoting the expression and secretion of inflammatory cues. Given the strong induction of *CXCL8* in our experiments and the established role of CXCL8 in neutrophil recruitment and early inflammatory amplification^28^, we next asked whether this pathway has clinical relevance in human KD and whether CXCL8 might serve as an early diagnostic biomarker to address this diagnostic gap. We therefore measured the serum CXCL8 concentration in the patient samples. KD patients displayed higher concentrations of CXCL8 in their sera compared with febrile controls, and importantly, this difference was already evident at the time of their first clinical visit (Figure 7A). This temporal pattern is consistent with CXCL8 acting early in disease initiation and supports its potential for early clinical discrimination between KD and other febrile illnesses.

**Figure 7.**
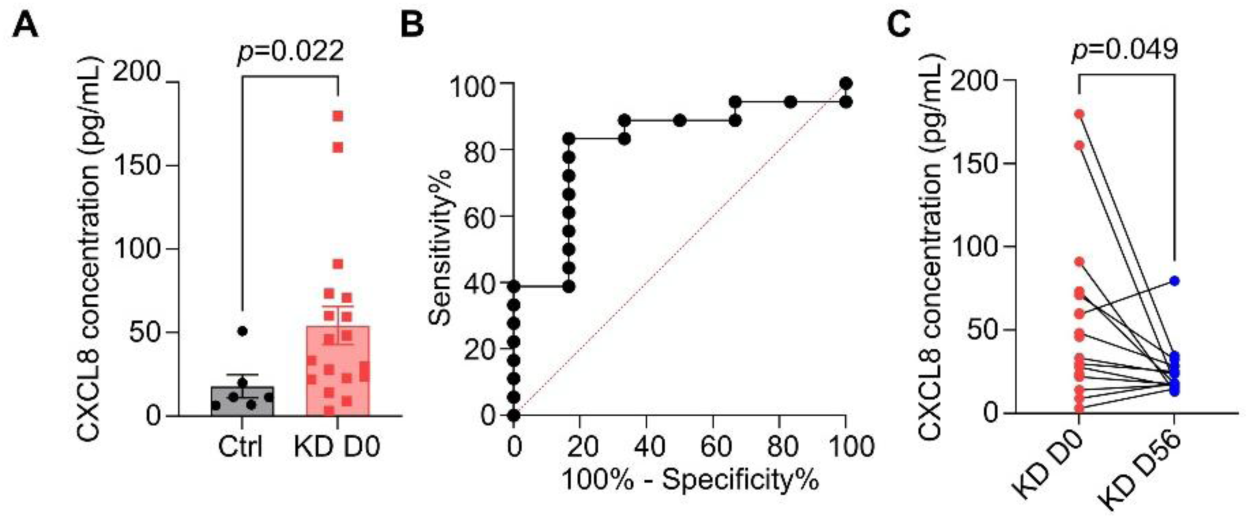
Serum CXCL8 is elevated in acute KD and demonstrates diagnostic utility at first clinical presentation. **A**, Serum CXCL8 concentrations measured by ELISA in febrile controls (Ctrl; n=6) and KD patients at day 0 (KD D0; n=18) of their initial ED visit. Serum CXCL8 is significantly elevated in KD patients prior to definitive diagnosis (Mann–Whitney U test; p=0.0224). **B**, Receiver operating characteristic (ROC) curve for serum CXCL8 in discriminating KD from febrile controls. At an optimal cut-off value of 20.91 pg/mL, CXCL8 achieved 83.33% sensitivity and 83.33% specificity. **C**, Longitudinal serum CXCL8 concentrations in paired KD patient samples collected at day 0 (KD D0; n=14) and day 56 (KD D56; n=14) after initial ED presentation. Serum CXCL8 declined significantly at 8-week follow-up, confirming its association with the acute inflammatory phase (Wilcoxon matched-pairs signed rank test; p=0.0494).

To quantify diagnostic performance, we generated a receiver operating characteristic (ROC) curve using serum CXCL8 concentrations. CXCL8 demonstrated diagnostic utility, achieving a sensitivity of 83.33% and a specificity of 83.33% at an optimal cut-off value of 20.91 pg/mL (Figure 7B). In addition, we longitudinally assessed serum CXCL8 levels in patients who returned for follow-up 56 days after their initial ED visit. At this time point, KD was clinically resolved, with no persistent fever. Serum CXCL8 levels were markedly reduced compared with those measured at the time of emergency presentation (Day 0), indicating that CXCL8 is specifically elevated during the acute phase of the disease (Figure 7C). These data connect the miR-10b-5p-MKI67/CBX5-CEBPA-CXCL8 axis observed in vascular endothelial cells to a clinically measurable inflammatory output, highlighting CXCL8 as both a downstream effector of endothelial reprogramming and a candidate early diagnostic biomarker for KD.

## Discussion

In this study, we discovered 14 upregulated miRNAs from buffy coats of acute KD patients and confirmed that miR-10b-5p is necessary for KD-like vasculitis *in vivo.* miR-10b-5p reprogrammed coronary artery endothelial and smooth muscle cells from proliferative/metabolic to an inflammatory state at the transcriptome level. miR-10b-5p also changed chromatin accessibility in the coronary endothelium, and CEBPA, a transcription factor, mediated *CXCL8* and *MMP10* activation. Acute KD patients exhibited higher levels of CXCL8 in their sera and this distinguished KD from other febrile illnesses.

Based on these findings, we suggest a novel miR-10b-5p-MKI67/CBX5-CEBPA-CXCL8 axis in KD vasculitis (Figure 8). First, miR-10b-5p is upregulated in the early stage of KD, and in the coronary artery endothelial cells, miR-10b-5p suppresses key genes related to cell cycle and metabolism, such as *MKI67*, *CBX5*, and *SPOCK1*. At the same time, CEBPA binds to *CXCL8* and *MMP10* promoter regions, upregulating these genes and shifting the coronary endothelium into an inflammatory state. Consequently, CXCL8 is secreted into the bloodstream and recruits circulating neutrophils to the tunica intima, initiating vascular inflammation.

**Figure 8.**
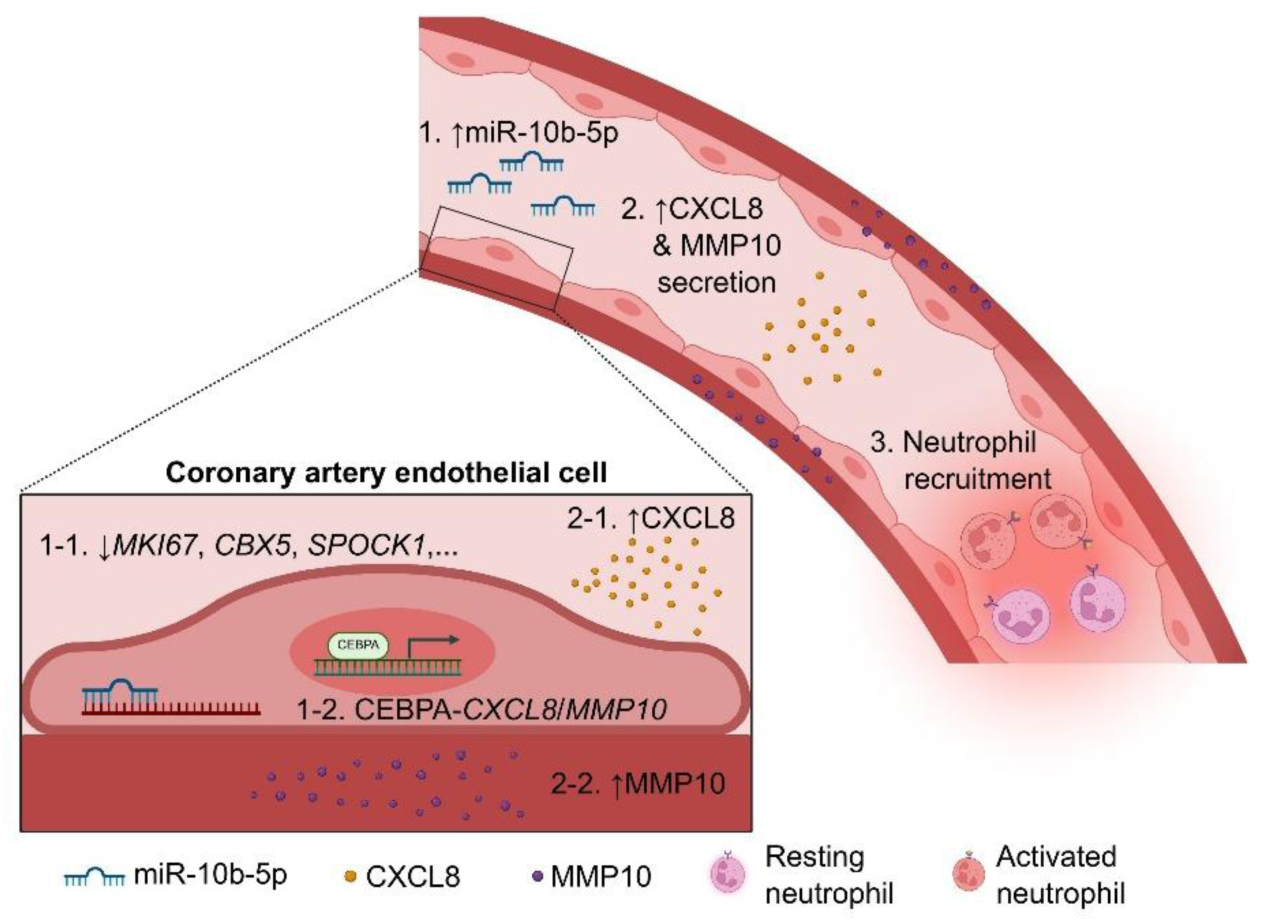
Working hypothesis: the miR-10b-5p–MKI67*/*CBX5–CEBPA–CXCL8 axis as a driver of endothelial inflammatory amplification in KD vasculitis. Schematic summarizing the proposed mechanism by which circulating miR-10b-5p contributes to vascular inflammation in KD. In the hyperacute/acute phase, elevated circulating miR-10b-5p enters coronary artery endothelial cells and suppresses proliferative/metabolic target genes, including *MKI67*, *CBX5*, and *SPOCK1* (step 1-1). Loss of these programs is accompanied by chromatin remodeling, increasing accessibility at inflammatory loci and enabling CEBPA to engage the *CXCL8* and *MMP10* promoters (step 1-2). Consequently, endothelial CXCL8 is secreted into the bloodstream (step 2-1), MMP10 is secreted into the tunica media (step 2-2), and eventually, circulating neutrophils are activated and recruited to the tunica intima (step 3), propagating vascular inflammation and contributing to KD-associated coronary artery injury. Created in BioRender.com.

This model aligns with the concept that endothelial cells represent an initiating site of immune activation in KD vasculitis. Vascular endothelial cells reside in the tunica intima, which first contacts with inflammatory molecules and cells in the blood^13^. During the acute and subacute stage of KD, activated monocytes, neutrophils, and natural killer (NK) cells are recruited and adhere to vascular endothelial cells^13^. These inflammatory cells release cytokines such as tumor necrosis factor (TNF) and interleukin-1β (IL-1β)^7^, which further trigger downstream pathological changes in endothelial cells, including cytokine production, reactive oxygen species (ROS) accumulation, and lipid oxidative stress, eventually developing coronary artery lesions^13^. In KD, therefore, endothelial cells function not merely as passive inflammatory targets, but as active amplifiers of vascular inflammation. Our findings extend this view by demonstrating that endothelial cells undergo active inflammatory reprogramming in KD, rather than serving as a passive target. Specifically, miR-10b-5p suppresses a proliferative/metabolic program (e.g., *MKI67*/*CBX5*) and activates an inflammatory mediator (e.g., CEBPA-driven *CXCL8* and *MMP10*), consistent with an endothelial amplifier that promotes neutrophil recruitment and vascular injury.

While endothelial activation may initiate immune cell recruitment, KD pathogenesis has also been viewed through the lens of vascular wall remodeling. Following injury, smooth muscle cells transition to myofibroblasts and produce matrix products^7,26^. These myofibroblasts proliferate and progressively injure the arterial wall^26^. Furthermore, vascular smooth muscle cells themselves can also dedifferentiate and proliferate to thicken and damage the vascular wall^27^. Together, these remodeling processes may represent key downstream structural outcomes that follow endothelial immune activation. Consistent with this remodeling program, our findings suggest that miR-10b-5p and subsequent endothelial reprogramming may provide an upstream trigger for these structural changes. Specifically, miR-10b-5p directly shifts endothelial cells and smooth muscle cells toward a pro-inflammatory state, which could in turn further promote smooth muscle cell phenotypic switching and progressive vascular wall thickening.

During the hyperacute/acute phase of KD, accumulating evidence supports a prominent role for innate immune activation, with neutrophils representing an early and functionally important component of the vascular inflammatory response^7,17,18^. Because endothelial cells constitute the first vascular interface with circulating leukocytes, endothelial induction of neutrophil-recruiting chemokines provides a plausible mechanism for the initiation of neutrophil-enriched inflammation observed early in KD vasculitis^17^.

Within this framework, CXCL8 provides a possible mechanistic bridge between endothelial reprogramming and KD immunopathology. CXCL8 is a neutrophil-attracting chemokine, and its induction in the coronary endothelium provides a potential mechanism linking miR-10b-5p-driven endothelial reprogramming to the early neutrophil recruitment. Our data provide direct evidence that miR-10b-5p suppresses *MKI67* and *CBX5*, and that CEBPA directly engages the *CXCL8* promoter to enable *CXCL8* transcription. While molecular steps linking *MKI67* suppression to activation of the CEBPA program require further definition, these results together support a miR-10b-5p-MKI67/CBX5-CEBPA-CXCL8 axis as a contributor to KD-associated vascular inflammation.

In addition to CXCL8, our data identify MMP10 as a CEBPA-linked output gene induced in the miR-10b-5p context. MMP10 belongs to the MMP family and has been implicated in inflammatory tissue remodeling through the degradation of ECM components, thereby facilitating leukocyte infiltration and structural injury within inflamed tissues^29,30^. From a vascular perspective, induction of MMP10 in the coronary endothelium is conceptually important because it suggests that the CEBPA program may not only recruit neutrophils via CXCL8 but also prime the vessel wall for matrix remodeling and barrier disruption, providing a mechanistic route toward downstream coronary artery lesion formation.

A notable finding is that the serum CXCL8 remains substantially elevated even at day 56 (Figure 7C), a time point at which acute inflammation is expected to have largely resolved in many KDs. This persistence raises the possibility that, even after apparent clinical resolution, a subclinical vascular inflammatory or remodeling program may remain active. This is consistent with the concept that vascular lesions can evolve beyond the hyperacute stage and that tissue remodeling processes may be prolonged. An additional, non-mutually exclusive explanation is that the inflammatory signal may be influenced by the stability and clearance kinetics of circulating miRNAs. Circulating miRNAs are known to be protected from degradation when packaged in extracellular vesicles or bound to proteins, enabling them to persist longer^31,32^. This would be particularly relevant if the dominant source of miRNAs includes platelets or activated leukocyte populations that can remain functionally altered after acute inflammation and can continue to contribute to extracellular miRNA pools^33,34^.

Several limitations should be considered. First, we were not able to define the cellular source of miR-10b-5p. Although circulating miRNAs can arise from multiple vascular and immune cell types, and may be exchanged via extracellular vesicles, the relative contributions of leukocytes, platelets, or other tissue compartments in KD remain incompletely resolved^35^.

Because we detected circulating miRNAs from the buffy coats, platelets and activated leukocytes are possible candidates and should be prioritized for follow-up.

Second, although the LCWE/CAWS model and the HCAEC model capture key inflammatory features, they cannot fully simulate human KD in infectious triggers or immune heterogeneity. In particular, KD is widely suspected to involve infectious or environmental triggers and exhibits substantial immune heterogeneity across patients^36^. These features are only partially modeled by the LCWE/CAWS challenge or endothelial-only *in vitro* systems. Incorporating additional human systems, such as patient-derived immune-endothelial co-cultures, may help capture this heterogeneity and refine translational inference.

Finally, although the CEBPA-*CXCL8* linkage is robust in the miR-10b-5p context, we cannot exclude additional parallel mechanisms. Inflammatory activation commonly involves multiple transcriptional regulators and chromatin modifiers^37^, and other transcription factors or epigenetic pathways could cooperate with, or act independently of, CEBPA to activate *CXCL8*. Thus, while our data support a miR-10b-5p-MKI67/CBX5-CEBPA-CXCL8 axis as one contributor to KD-associated vascular inflammation, delineating the full regulatory network will require systematic perturbation and chromatin-resolved mapping in future studies.

For future directions, we should investigate serum CXCL8 levels in a prospective cohort and evaluate the IVIG response and coronary artery lesions using these values. We also propose a single-cell multiome analysis combined with extracellular vesicle profiling to clarify the source of miR-10b-5p and downstream regulatory pathways.

Together, our study positions miR-10b-5p as an upstream regulator that links endothelial chromatin/transcriptional reprogramming to neutrophil-recruiting inflammation in KD, with CXCL8 as a key CEBPA-driven output. This miR-10b-5p-MKI67/CBX5-CEBPA-CXCL8 axis provides a framework that connects early endothelial immune activation to downstream vascular injury and remodeling, and it nominates CXCL8 as a clinically relevant biomarker in acute KD.

## Acknowledgement

All authors thank to Prof. Eun Jeong Won MD, PhD (University of Ulsan College of Medicine) for her assistant in the preparation of CAWS.

## Author Contributions

Drs G. H. Eom and S. Yoon designed, supervised, and financially supported the studies. Dr. Y. K. Cho designed the study and enrolled patients. Dr. G. Kang financially supported the studies and wrote the manuscript. Ms S. Park analyzed bioinformatic data and wrote the manuscript. Ms M. Kim and Ms B. Lim performed the *in vivo* and *in vitro* experiments. Drs H. Yang and S. Jung collected the human specimen. Drs M. H. Kim and W-J. Park performed miRNA transduction and scRNA-seq. Dr. Y-K. Kim analysed whole miRNA-seq data. All authors have read and approved the article.

